# Vaccine-elicited and naturally elicited antibodies differ in their recognition of the HIV-1 fusion peptide

**DOI:** 10.1101/2024.06.18.599578

**Authors:** Mateo Reveiz, Kai Xu, Myungjin Lee, Shuishu Wang, Adam S. Olia, Darcy R. Harris, Kevin Liu, Tracy Liu, Andrew J. Schaub, Tyler Stephens, Yiran Wang, Baoshan Zhang, Rick Huang, Yaroslav Tsybovsky, Peter D. Kwong, Reda Rawi

## Abstract

Broadly neutralizing antibodies have been proposed as templates for HIV-vaccine design, but it has been unclear how similar vaccine-elicited antibodies are to their naturally elicited templates. To provide insight, here we compare the recognition of naturally elicited and vaccine-elicited antibodies targeting the HIV-1-fusion peptide, which comprises envelope (Env) residues 512-526, with the most common sequence being AVGIGAVFLGFLGAA. Naturally elicited antibodies bound peptides with negative-charge substitutions around residues 517-520 substantially better than the most common sequence, despite these substitutions rarely appearing in HIV; by contrast, vaccine-elicited antibodies were less tolerant of sequence variation, with no substitution of residues 512-516 showing increased binding. Molecular dynamics analysis and cryo-EM structure of the naturally elicited ACS202 antibody in complex with HIV-Env trimer with A517E suggested enhanced binding to result from electrostatic interactions with positively charged antibody residues. Overall, vaccine-elicited antibodies appeared to be more fully optimized to bind the most common fusion peptide sequence.

**HIGHLIGHTS:**

Peptide substitution scan reveals naturally elicited antibodies against fusion peptide (FP) can bind select non-canonical FP sequences with high affinity.
Peptide substitution scan data for FP antibodies correlates significantly with their differential selection indicating variation in binding relates to neutralization tolerance.
Structure and energetic analysis of naturally elicited ACS202 with HIV-Env trimer reveals basis for improved recognition of A517E mutant.
Atomic level interactions from MD simulation analysis corroborate trends observed with peptide substitutions.
verall, peptide substitution scans reveal vaccine-elicited antibodies against FP to be less permissive to FP-sequence variability than naturally elicited antibodies.

## INTRODUCTION

Two general strategies – lineage-based and epitope-based – have been proposed by which to use broadly neutralizing antibodies as templates for HIV vaccine development ^1–3^. In lineage-based strategies, antibodies of the same reproducible class are elicited by priming with immunogens that preferentially bind to initial recombinants, followed by additional boost to ensure appropriate lineage maturation, with a final polishing step to expand the expression of the vaccine-elicited broadly neutralizing antibodies ^4–7^. This approach is achieving substantial success in knock-in mice models ^8,9^ and in the initial stage of priming in human clinical trials ^10^. In epitope-based strategies, sites of vulnerability on the HIV-1 Env trimer are defined by antibodies with broad and potent neutralization, and immunogens focus the immune response to specific sites, and success has been achieved through diverse approaches involving (i) priming with Env trimers with *N*-linked glycans proximal to the target site removed to enhance local immunogenicity ^11,12^ or (ii) priming with peptides or nanoparticle-based scaffolds that antigenically mimic the target site ^13–16^.

Success with epitope-based design has been particularly noteworthy at the fusion peptide-site of vulnerability, with broadly neutralizing responses achieved in mice, guinea pigs, and non-human primates ^17–19^. This breakthrough has allowed us to analyze the degree of similarity between the vaccine-elicited antibodies and the naturally elicited ones.

In this study, we measured the impact of single point mutations on the fusion peptide, residues 512-526, using the most common fusion peptide sequence (_512_AVGIGAVFLGFLGAA_526_; FP15) on the binding of eight vaccine-elicited antibodies and three naturally elicited antibodies. The analysis was based on a PEPperMAP^®^ epitope substitution scan of all amino acid positions with the 20 canonical amino acids ^20^. The naturally elicited antibodies included three human antibodies: PGT151 (isolated from an elite neutralizer infected with a clade C virus; 60% breadth on 208 panel) ^21^, N123-VRC34.01 (isolated from a donor infected with a clade B virus; 50% breadth) ^22^, and ACS202 (isolated from an individual infected with a subtype B virus; 26% breadth) ^23^. For vaccine-elicited antibodies, we selected three murine antibodies ^15^, vFP1.01, vFP5.01 and vFP16.02 as well as five macaque antibodies ^18^, 0PV-a.01, 0PV-b.01, 0PV-c.01, DF1W-a.01, and DFPH-a.01. Select substitutions were assessed for altered affinity in the context of prefusion-closed Env trimers, the cryo-EM structure of naturally elicited antibody ACS202 was determined in complex with one of the substitutions with most enhanced affinity, and molecular dynamics analysis was used to provide mechanistic insight into affinity improvement. Overall, naturally elicited antibodies showed substantially higher affinity to fusion peptide variants, most of which were not commonly present in HIV-1, whereas the vaccine-elicited antibodies appeared to be optimized in their recognition of the most common sequence variant of the fusion peptide.

## RESULTS

### Peptide substitution scans for naturally elicited fusion peptide-directed HIV-1 Env neutralizing antibodies reveal improved binding to non-canonical fusion peptide sequences

Naturally elicited, fusion peptide-directed, broadly neutralizing antibodies, VRC34.01, PGT151, and ACS202, assume multiple angles of approach when binding the HIV-1 trimer proximal to the FP site (**Figure 1A**). To provide insight into sequence requirements for binding, epitope substitution scans were performed with PEPperPRINT FP15 peptide microarrays (**Figure 1B**). These scans reveal that all three naturally-elicited human antibodies can bind diverse FP sequences.

**Figure 1.**
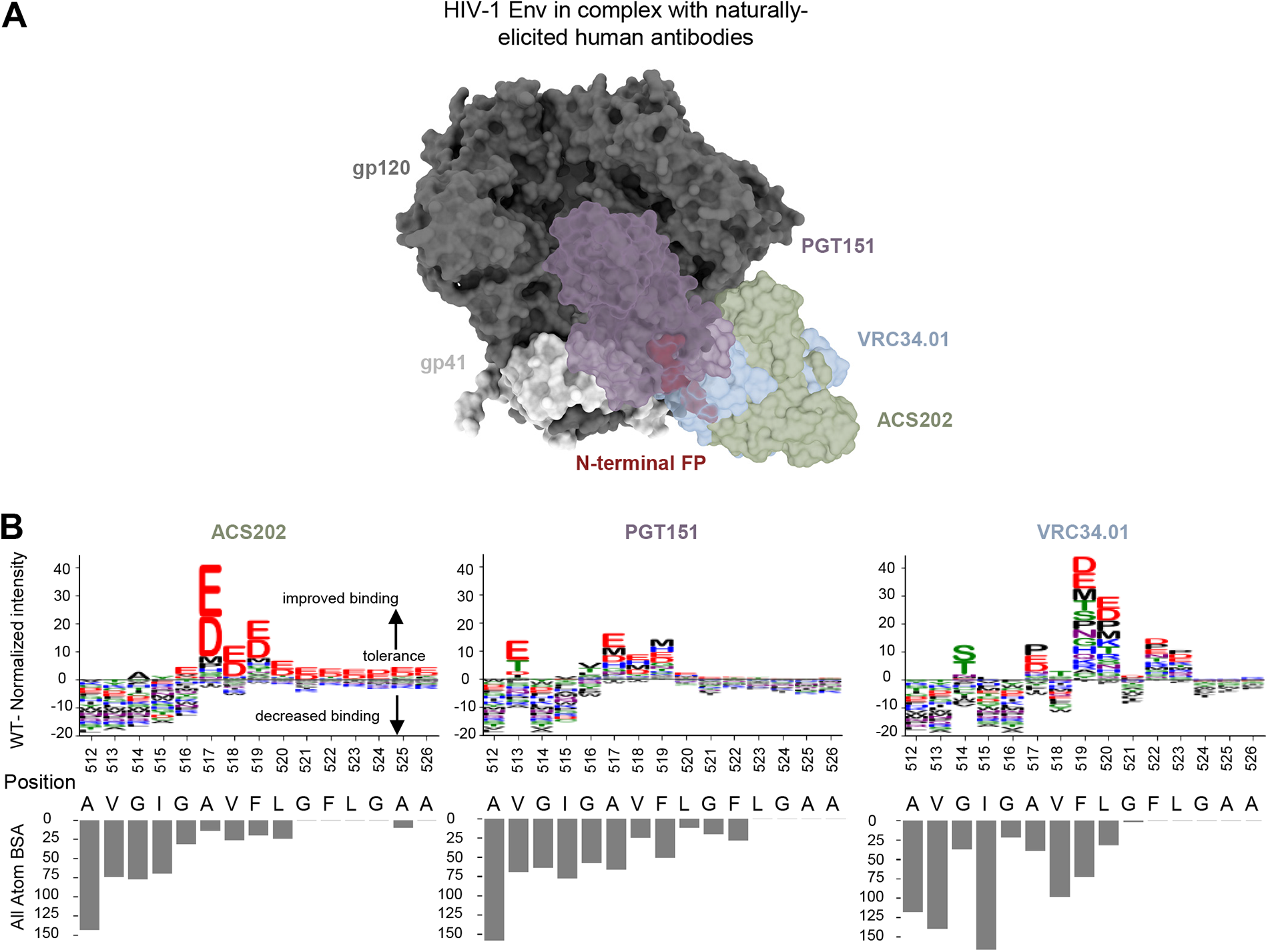
Diverse fusion peptide sequences are target sites for naturally elicited antibodies, as revealed by peptide-substitution analysis. (A) Structural representation of the HIV-1 Env trimer in complex with naturally elicited FP-targeted antibodies. (B) Logo plots with peptide substitution scan data from PEPperPRINT with all atom BSA bar plots. The height of each amino acid mutation corresponds to the relative improvement over WT, with positive values improving over WT, negative values decreasing binding over WT.

ACS202 tolerated single substitutions at multiple position (514-526). ACS202 analysis displayed a conserved N-terminal core A_512_VGIG_516_, a highly variable A_517_VF_519_ region with a preference for the negatively charged residues Asp and Glu with a 10-fold binding intensity increase with respect to wildtype (WT), as well as the hydrophobic residues Pro and Met to a much lower degree. The C-terminal region L_520_GFLGAA_526_ tolerated most mutations and displayed a consistent preference for negatively charged residues over WT. Overall conserved regions correlated well with all-atom buried surface area (BSA).

PGT151 on the other hand showed more moderate increases in normalized intensities at a few selected positions 513, 516-520. The N-terminal core A_512_VGIG_516_ remained conserved except for position 513 where most residues are more tolerated than with ACS202 and Glu, Thr, Asp and Ile are preferred over wildtype. The central region is once again highly variable with position 516 preferring Val and Thr. Most substitutions in the A_517_VF_519_ region improve over WT. The C-terminal region L_520_GFLGAA_526_ tolerated most mutations but did not show the strong improvement with negatively charged residues as shown in ACS202.

The third naturally elicited antibody N123-VRC34.01 (referred to as VRC34.01) showed strong conservation and preference for WT residues at positions 512-513, 515-516. Mutations at position 514 were slightly more tolerated, with a strong preference for Ser and Thr residues and a lower BSA than its neighbors. Similarly, mutations in position 517 allowed for certain flexibility with strong improvement over WT with Pro, Glu and Asp mutations. Positions 519 and 520 not only tolerated all single amino acid substitutions; they also all improved over wild type.

Overall, naturally-elicited antibodies displayed a conserved A_512_VGIG_516_ region with some exceptions while tolerating most substitutions in the C-terminal region. Multiple non-canonical mutants significantly improved binding over wildtype, with a strong preference for charged residues which are not usually found in actual HIV sequences.

### Peptide substitution scans for vaccine-elicited FP-directed HIV-1 Env neutralizing antibodies reveal lower tolerance to FP variability in the N-terminal region

While vaccine-elicited FP-directed murine NAbs vFP16.02 and vFP1.01 assume similar angles of approach when binding the HIV-1 trimer proximal to the FP site (**Figure 2A**), Neutralizing antibodies elicited in NHP assume multiple angles of approach when binding the HIV-1 fusion peptide (**Figure 1A**). Epitope substitution scans were also performed with PEPperPRINT FP15 peptide microarrays (**Figure 2B**). Overall, these scans revealed a much lower tolerance for single mutations in the N-terminal core A_512_VGIG_516_ suggesting the wildtype residues remain key for binding. The only exception is 0PV-b.01 which shows reduced antibody response above the noise level of the assay. There was clear promiscuity in the A_517_VFLGFLGAA_526_ region without significantly improved binders over WT for DF1W-a.01, DFPH-a.01, 0PV-b.01 and vFP1.01. The V_518_FL_520_ region showed moderate improvement over WT for 0PV-a.01 and vFP16.02. Interestingly 0PV-c.01 showed great improvement at position 519 and 520 but did not tolerate mutations at position 518.

**Figure 2.**
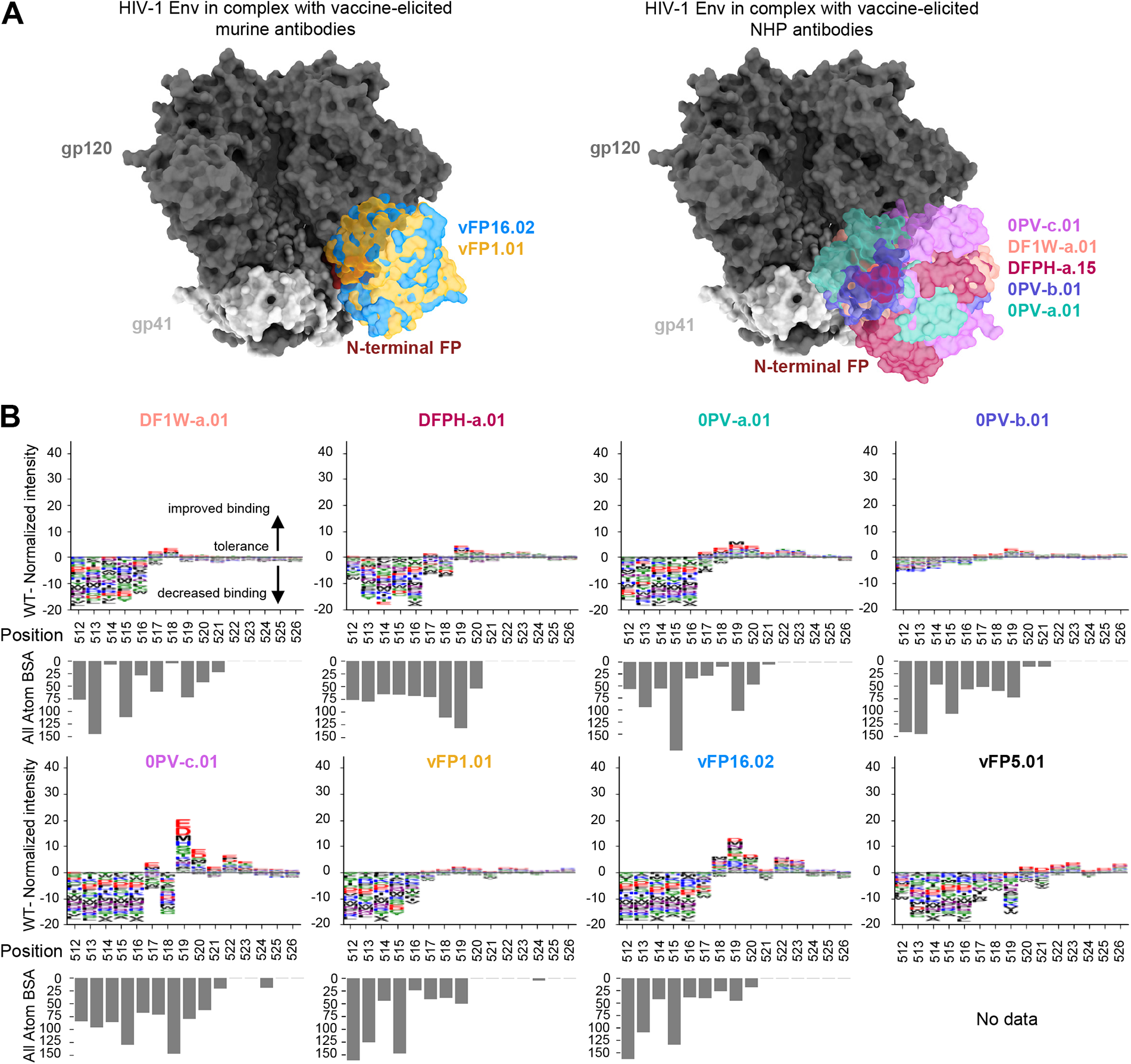
Peptide-substitution analysis reveals vaccine-elicited antibodies targeting FP site of vulnerability are less permissive to FP variation. (A) Overall structure context of vaccine elicited antibodies in complex with HIV-1 Env trimer. The structure from DFPH-a.15 is displayed for DFPH-a.01. (B) Logo plots with peptide substitution scan data from PEPperPRINT with all atom BSA bar plots. The height of each amino acid mutation corresponds to the relative improvement over WT, with positive values improving over WT, negative values decreasing binding over WT. DFPH-a.01 BSA

Overall, the vaccine-elicited antibodies appeared to be more optimized for the most common FP sequence in the N-terminal region (residues 512 through 516). In the 517-520 region, residue alterations did not substantially affect binding, although it was interesting to see that some of the same substitution observed in substantially improve affinity with naturally elicited antibodies also positively impacted the affinity for the vaccine-elicited antibodies, just to a much lower degree. And in the C-terminal region (521-526), there was generally little interaction with antibodies, and most substitutions had little impact.

### Statistically significant difference in diversity of FP sequences bound by vaccine elicited and naturally elicited antibodies

To quantitively compare naturally and vaccine elicited antibodies, the number of PEPperPRINT binding intensity values for the 19 mutants *I_mut_* higher than the wildtype value *l_wt_* were counted after grouping by FP position and antibody (**Figure 3A top panel**). Position 512 was absolutely conserved for both vaccine and naturally elicited antibodies. For positions 513-516, almost no mutations improved binding to vaccine elicited antibodies whereas 10-20% of mutants improved binding to naturally elicited antibodies. Mutations in the 517-520 region display very high levels of improvement (around 50%) for both types of antibodies. The magnitude of the responses *M* was also assessed by averaging the wildtype-normalized PEPperPRINT binding intensity values *I* per FP position *i* across all 19 amino acid substitutions, as shown by 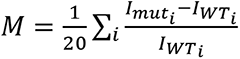 (Figure 3A bottom panel). Here significant differences were observed between naturally and vaccine elicited antibodies for all positions except for positions 512 which was highly conserved, as well as position 520.

**Figure 3.**
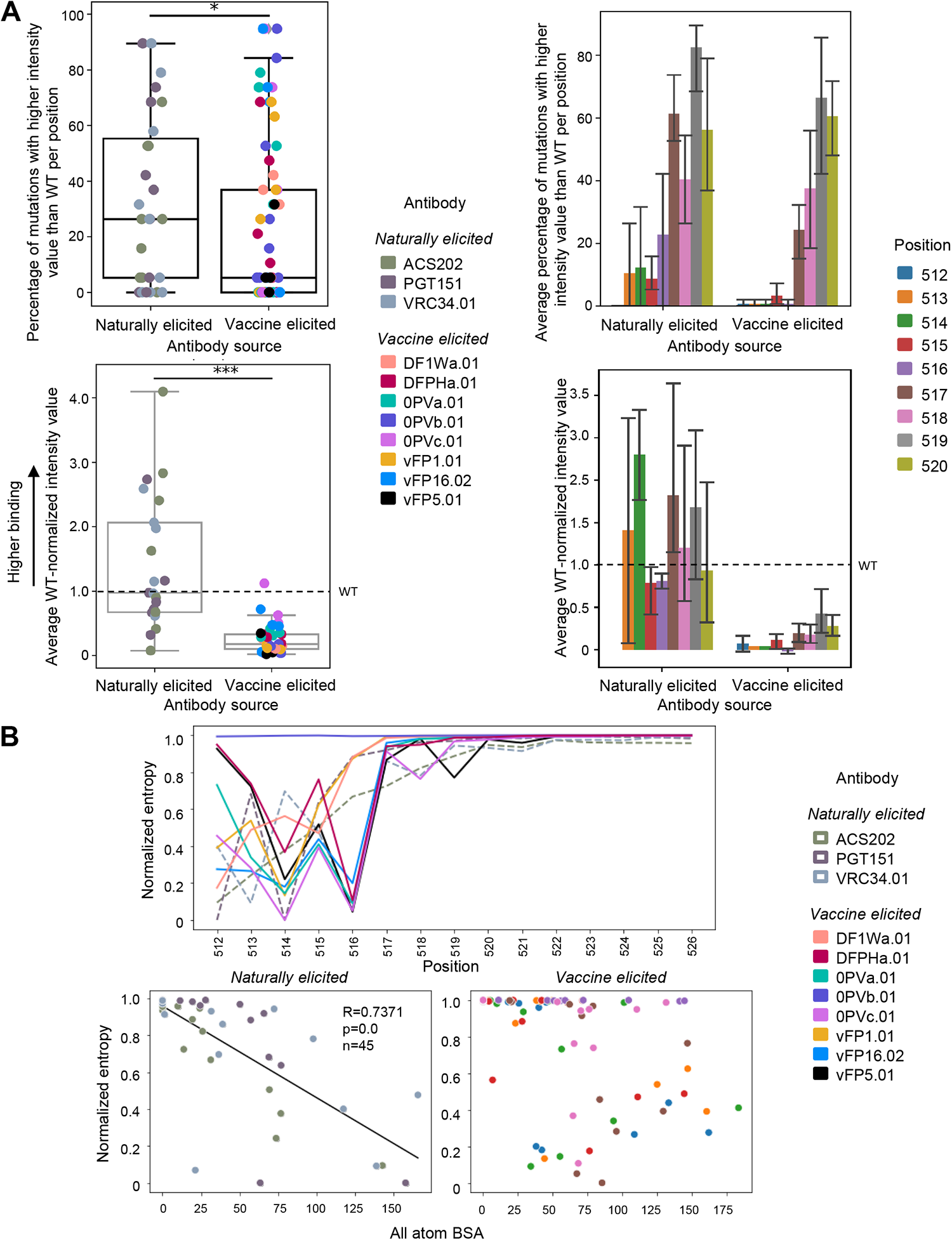
Quantification of FP recognition reveals naturally-elicited antibodies bind significantly more diverse FP sequences than those that are vaccine-elicited. (A) Top panels: Count analysis: Number of mutation with higher PEPperPRINT intensity value than WT for positions 512-520. On the left, each point corresponds to a particular position and antibody for n=72 vaccine elicited and n=27 for naturally elicited antibodies. On the right: values are averaged across antibodies for a given position. Bottom panels: Magnitude analysis: Average PEPperPRINT intensity value for entries higher than WT for positions 512-520. Error bars correspond to 95% confidence interval. (B) The intensity values for each position are normalized and used to compute per-position entropies with base=20 to normalize. Entropies for both naturally and vaccine elicited antibodies are correlated with all atom BSA.

The distribution of PEPperPRINT intensities was used to compute the entropy for a given position (**Figure 3B**). Positions 518 to 526 displayed the maximum entropy for both vaccine and naturally elicited antibodies, suggesting a uniform distribution over all possible mutations. Naturally elicited antibodies displayed high correlation between entropy and all-atom BSA, with low entropy positions having the highest BSA values (R=0.73, P<1e-5). In other words, exposed residue positions showed little preference to mutations while buried positions had specific preferences for certain mutations. However, some clear exceptions exist; positions such as 517 in complex with ACS202 can be easily mutated while having very low BSA of 10 A^2^, whereas 519 in VRC34 has a high BSA of 75 A^2^ and tolerated most substitutions. Surprisingly, vaccine elicited antibodies did not seem to follow such a trend, with very different clusters for different positions. The strong preference to certain residues regardless of the antibody contacts might relate to the initial priming during vaccination.

### PEPperPRINT binding data for FP antibodies correlates significantly with other established metrics including differential selection and binding in trimer context by Octet

BG505 trimers with specific mutations of interest spanning diverse PEPperPRINT responses were created to check if peptide binding would correlate with its corresponding trimer context (**Figure S2A**). In particular, mutation A517E was selected since it strongly improves binding across all naturally elicited antibodies while maintaining a small change in vaccine elicited antibodies. Mutation G514S was also selected since it strongly and uniquely improves binding with VRC34 antibody. The binding response was measured in triplicates for the mutants and subtracted by the wildtype antibodies to compute Δ Binding responses. Overall, mutation 517E improved binding, except for VRC34 and DF1W-a.01. The G514S mutation improve VRC34 as expected, but also increased binding to DF1W-a.0, contradicting PEPperPRINT prediction in peptide context. PEPperPRINT peptide binding values show a medium correlation (R=0.6828, p=0.00179) with Octet binding in trimer context (**Figure S2A, S2B**).

In addition to trimer context, PEPperPRINT binding values correlated with differential selection metrics ^24,25^, suggesting that variation in binding somewhat relates to neutralization tolerance (**Figure S1**).

### Cryo-EM structure of ACS202 in complex with BG505 Env trimer with A517E explains enhanced affinity of A517E substitution

To elucidate the structural mechanism for the improvement of antibody binding to the A517E mutant, we determined a cryo-EM structure of ACS202 Fab in complex with BG505_RnS_7mut_A517E, a prefusion-stabilized Env trimer ^26^ with the alanine at position 517 mutated to glutamate. We obtained a 3D-reconstruction map at 2.30 Å from 635,332 particles applying C3 symmetry (**Figure S3; Table S1**). Local resolution estimation indicated that the antibody bound with substantial flexibility, with the Fab having substantially lower resolution than Env trimer (**Figure S3 panel E**). However, the antibody-antigen interface was well defined in the density map and showed that ACS202 interacted with the N-terminal segment of the fusion peptide through β-strand interactions from its third complementarity determining region of heavy chain (CDR H3) (**Figure 4A**), similar to that observed for the previously determined ACS202-AMC011 complex ^27^. The electron density was relatively weak around residue 517, especially around the glutamate side chain (**Figure 4B**), indicating conformational flexibility. Structural alignment with the ACS202-AMC011 complex (PDB: 6nc2) ^27^ indicated that A517E mutation likely induced local conformational flexibility. AMC011 has the N-terminal 15 residues of fusion peptide identical to that in BG505, with alanine at residue 517. The two structures were aligned by one heavy chain variable domain of ACS202, with a rmsd of 0.677 Å for 126 aligned Cα atoms (**Figure 4C**). Both heavy and light chains, as well as most of the protomer (gp41 and gp120) interacting with the aligned antibody, were well aligned. However, residues 517-518 had substantial shifts, with main chain atoms between the two structures having up to ∼4 Å apart. The conformational flexibility around E517 could potentially allow the nearby R100f side chain on CDR H3 to have more favorable interactions with the carbonyl groups of residues 516-517 or to have dynamic charge-charge interactions with E517 side chain, although the refined cryo-EM structure exhibited a distance of 7.8 Å between the two oppositely charged side chains. Structural constraints from the rest of the trimer-antibody interactions likely limits the interactions between the two side chains. This is consistent with our observation that PEPperPRINT analysis of amino acid substitutions in the fusion peptide, which was not constrained by trimer interactions, exhibited substantial enhanced binding to ACS202 with glutamate or aspartate at residue 517 (**Figure 1B**), whereas when measured by Octet in the context of trimer, A517E mutation exhibited a much smaller increase in binding to ACS202 (**Figure S2**).

**Figure 4.**
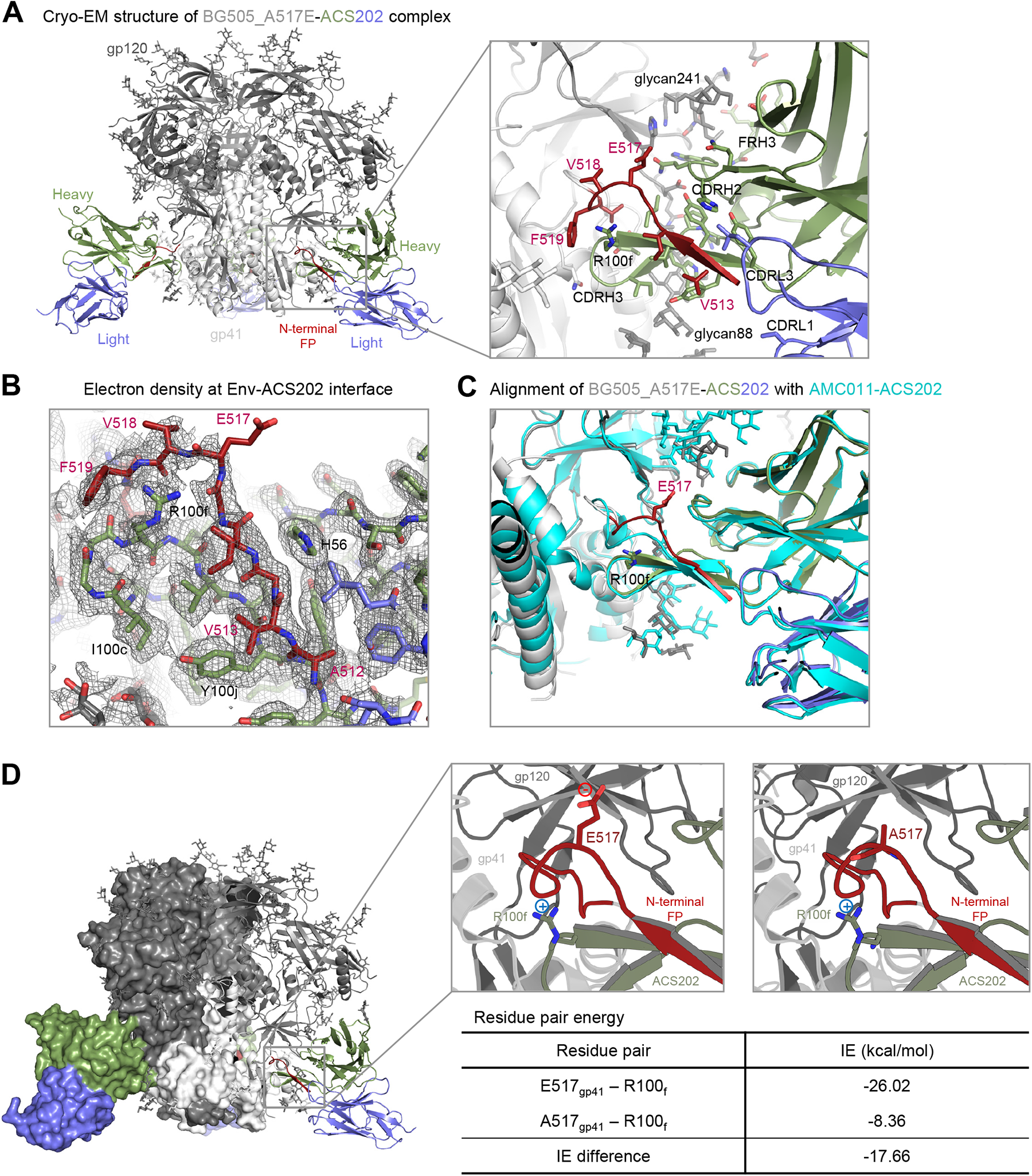
Cryo-EM structure of ACS202 in complex with BG505 Env trimer with A517E explains enhanced affinity of A517E substitution. (A) Cryo-EM structure of BG505_A517E in complex with ACS202. (B) Electron density at the trimer-antibody interface, contoured at 4 σ. The map was from final cryoSPARC non-uniformed refinement, sharpened, at a nominal resolution of 2.3 Å. (C) Structural alignment of BG505_A517E-ACS202 and AMC011-ACS202 complexes. The structures were aligned by the antibody heavy chain shown. (D) Key residue pair in the BG505_A517E-ACS202 complex contributed to enhanced binding affinity.

To confirm the dynamic interactions between the side chains of E517 at the fusion peptide and R100f of CDR H3, we conducted an analysis of the energetic interactions between these residue pairs using the trajectory obtained from molecular dynamics simulations (**Figure 4D**). Pairwise residue energy was calculated between R100f and residue 517 as either alanine or glutamate, and a decrease of 17.66 kcal/mol was observed for the A517E mutation, indicating that there were favorable dynamic interactions between residues R100f and E517.

Overall, the cryo-EM structure of BG505_A517E in complex with ACS202 revealed that, in the context of trimer-antibody binding interactions, the fusion peptide was unable to adopt a stable conformation that brought the side chains of E517 and R100f close enough for salt-bridge interaction. However, the A517E mutation induced conformational flexibility around residue 517 in the fusion peptide of the trimer-antibody complex, allowing for favorable interaction between residues E517 and R100f, contributing to the observed improvement in affinity, although in a smaller scale than that observed for the free fusion peptide binding to the antibody ACS202. This favorable dynamic interaction could be between the two side chains or between the R100f side chain and the main-chain carbonyl groups of residues 517-518.

### Atomic level interactions from MD simulation analysis corroborate trends observed with PEPperPRINT intensity values

To further understand the structural mechanisms explaining the improved binding to naturally elicited antibodies, we selected two mutations and analyzed the trajectories resulting from MD simulations. For the mutant selection, we first calculated the delta PEPperPRINT intensities Δ*I* independently for the vaccine elicited antibodies VRC34 and PGT151 by subtracting the average naturally elicited intensity values for each position and mutation (**Figure 5A, S4A**). The highest Δ*I* values corresponding to the highest expected differences between vaccine and naturally elicited antibodies were G514S for VRC34 and V513E for PGT151. The wildtype and mutated versions of the FP were generated in complex with the corresponding naturally elicited antibodies and the vaccine elicited antibodies 0PVc.01 and vFP16.02, which also showed the highest Δ*I* values (8 total complex structures). The structures were used to run 3 independent sets of MD simulations. The resulting trajectories were analyzed by running PCA on C-alpha atoms and clustered on the 2D projections using the meanshift algorithm ^28^. The centroid of each cluster was computed and the closest MD frame to the centroid by euclidean distance was used as a representative frame (**Figure S4B**). These frames were then used to compute the average pairwise energies of centroid conformations. The average pairwise energies were used to compare between the wildtype and mutated forms by subtracting the pairwise energies resulting in Δ*E* maps (**Figure 5C, Figure S5**).

**Figure 5.**
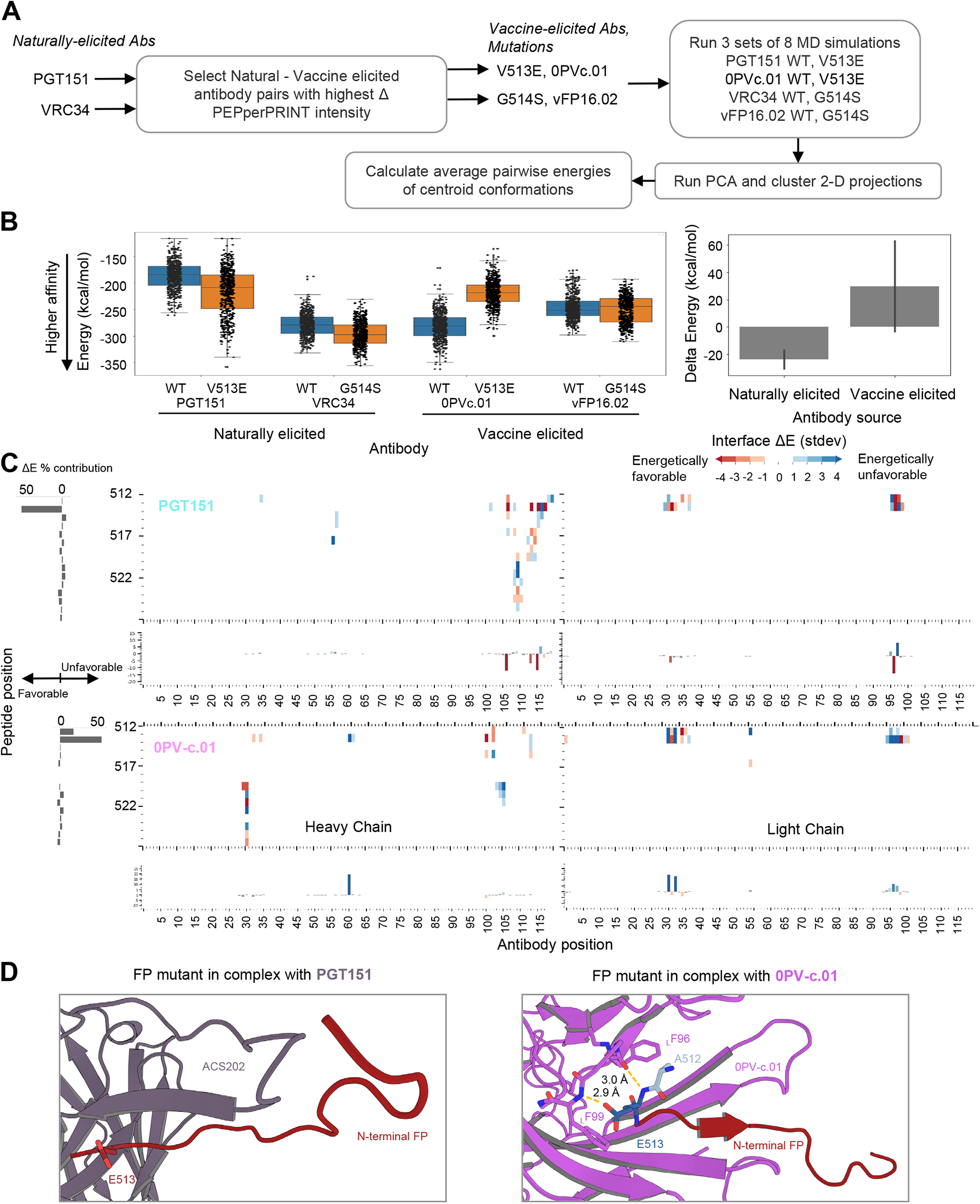
Atomic level interactions from MD simulation analysis corroborate trends observed by peptide-substitution analysis. (A) Diagram for *in silico* structural analysis pipeline of FP mutants. (B) Overall energies suggest mutations on naturally elicited antibodies improve binding whereas vaccine elicited antibodies do not tolerate those mutations as well. (C) Pairwise energy analysis reveals specific interactions responsible for the binding differences observed in PEPperPRINT. Numbering of residues is sequential and not in kabat format. (D) MD structures of FP mutants in complex with antibodies. FP residues are color coded by the energy contributions from C, with red indicating more favorable energetic interactions and blue indicating unfavorable.

From the PEPperPRINT Δ*I* values, we expected an increased binding for naturally elicited antibodies and a decrease in mutated residues for vaccine elicited antibodies. The interaction energy between the antibody and FP complex was computed using NAMD. The expected trend is observed for the naturally elicited antibodies PGT151 and VRC34, where the 513E and 514S mutants show a slight decrease in energy from an average of -186 to -217 kcal/mol and from -279 to -296 kcal/mol respectively (**Figure 5B**). The most dramatic and consistent change across all antibodies is observed for 0PVc.01 513E mutation with an energy increase from -282 to -219 kcal/mol. The vPF16.02 shows a negligeable decrease with mutation 514S mutation from -246 to -250 kcal/mol. Overall, the naturally elicited antibodies show the expected trends of lowering the interaction energies although with significant overlap across the full trajectories.

## DISCUSSION

The vaccine elicitation of antibodies capable of broad and potent neutralization of HIV-1 is in a nascent stage. Antibodies isolated applying lineage-based vaccine strategies have achieve substantial breadth in knock-in mice (refs), while antibodies from epitope-based strategies have achieved over 50% breadth only against the fusion peptide-site of vulnerability (Kong et al., 2019). Because antibodies elicited by the latter strategy derived from different antibody classes than the template naturally elicited antibodies, we felt it would be important to assess the similarity by which naturally elicited and vaccine-elicited antibodies recognized fusion peptide.

Overall vaccine-elicited antibodies showed decreased tolerance to sequence variation, a general preference for the most common FP sequence and inconsistent correlations between full-atom buried surface area (BSA) and PEPperPRINT binding entropy for each position. On the other hand, naturally elicited antibodies showed on average a consistent binding to FP, with some mutations strongly improving binding over the consensus FP, as well as consistent BSA and entropy. In addition, the trajectories from MD simulations for selected FP mutations suggests lower interaction energies with naturally elicited antibodies, and higher interaction energies to vaccine-elicited antibodies, consistent with the observed binding values from the substitution scan analysis. Although statistically significant differences were found between naturally and vaccine elicited antibodies, it is still possible that the observed differences are caused by the different origins of these antibodies (human versus mice/macaque).

It will be interesting to see if the revealed differences in fusion peptide recognition can be used to improve the breath and potency of vaccine-elicited antibodies. It was interesting to observe that while naturally elicited antibodies often have superior potency – they were not fully optimized in terms of binding to the most commonly occurring FP15 sequence. The fact that similar changes at residues 517-520 positively impacted the recognition of naturally elicited antibodies (and to a lesser degree the vaccine-elicited ones) suggests that antibodies have not fully optimized their electrostatics interactions. The much lower enhancement in affinity noted with the vaccine-elicited antibodies does suggest, however, that they were optimized to recognize exposed N-terminus of FP, perhaps a consequence of their priming with the most common fusion peptide sequences.

## Acknowledgments

We thank Allison Zeher, AJ Morton, and Zabrina Lang for data collection on the Titan Krios operated by NIH IRP cryo-EM consortium.

## Author contributions

M.R., K.X., and R.R. performed analyses and conceptualized project; M.L. performed MD analysis; S.W. determined cryoEM structure of BG505_A517E in complex with ACS202 Fab; A.S.O. and D.R.H. produced BG505-A517E Env trimer and performed Octet binding; T.L. and B.Z. produced ACS202 Fab; K.L. and Y.Y. contributed to reagent preparation; A.J.S. assisted with Env trimer visualizations; T.S. and Y.T. prepared grids; R.H. collected cryoEM data; and P.D.K. conceptualized project. M.R., S.W., P.D.K., and R.R. wrote the manuscript with all authors providing comments and feedback.

## MATERIALS AND METHODS

### Preparation of antibodies

In this study, a total of three human, three murine, and five macaque antibodies were generated. The DNA encoding the variable domains of the heavy and light chains for each antibody was codon optimized, synthesized, and cloned into specific VRC8400-based mammalian expression vectors. These vectors contained species-matched constant domains. Plasmids carrying the individual antibody heavy and light chain sequences were co-transfected into Expi293 cells using Turbo293 transfection reagent following the manufacturer’s protocol. The transfected cells were then incubated in shaker incubators at 120 rpm, 37°C, and 9% CO_2_. On the second day, AbBooster medium was added to the cell cultures at one-tenth of the culture volume, and the incubation temperature was lowered to 33°C while maintaining the other conditions. The cell cultures were further incubated for an additional 5 days. After 6 days post-transfection, the cell culture supernatants were harvested. Antibodies were purified from the supernatants using protein A chromatography. The purification process involved a PBS wash followed by elution with a low pH glycine buffer. The eluted antibodies were immediately neutralized using 10% volume of 1 M Tris buffer at pH 8.0.

### PEPperMAP® epitope substitution scan

The substitution scan analysis was performed as described in PEPperPRINT methods. Pre-staining of an AVGIGAVFLGFLGAA peptide microarray copy was done with the secondary goat anti-human IgG (Fc) DyLight680 antibody and the control antibody in incubation buffer to investigate background interactions with the variants of wild type peptide AVGIGAVFLGFLGAA that could interfere with the main assays. Subsequent incubation of further AVGIGAVFLGFLGAA peptide microarray copies with the synthetic human antibodies at concentrations of 1 µg/ml, 10 µg/ml and 100 µg/ml in incubation buffer was followed by staining with secondary and control antibodies as well as by microarray read-out at scanning intensities of 5/7 or 7/7 (red/green). The additional HA control peptides framing the peptide microarrays were simultaneously stained as internal quality control to confirm the assay quality and the peptide microarray integrity.

Quantification of spot intensities and peptide annotation were based on the 16-bit gray scale tiff files at a scanning intensities of 5/7 or 7/7 that exhibit a higher dynamic range than the 24-bit colorized tiff files included in this report; microarray image analysis was done with PepSlide^®^ Analyzer. A software algorithm breaks down fluorescence intensities of each spot into raw, foreground and background signal (see “Raw Data” tabs), and calculates averaged median foreground intensities and spot-to-spot deviations of spot triplicates (see “Data Summary” tabs). Based on averaged median foreground intensities, intensity maps were generated and interactions in the peptide maps highlighted by an intensity color code with red for high and white for low spot intensities.

Correlations with differential selection were performed by extracting the relevant metrics from the Dingens and Doud manuscripts and corresponding GitHub repositories https://github.com/jbloomlab/MAP_NHP_FP_Abs/tree/master ^24,25^.

### HIV-1 Env trimer production

Soluble HIV-1 Env trimers were expressed and purified as previously described ^29^. Briefly, plasmid encoding an N-terminal single chain Fc tag and HRV 3C cleavage site followed by either BG505-RnS or its mutants were transfected into HEK293F Freestyle cells. The protein was expressed for 6 days at 37°C, after which the supernatant was cleared by centrifugation and filtration. The HIV-1 trimers were purified by affinity chromatography with Protein A resin, and the Env trimer liberated from the tag by cleavage with HRV 3C. The eluted Env trimer was further purified by gel filtration on a Superdex S-200 column, equilibrated in PBS. The proteins were concentrated to 1mg/ml, flash frozen with 10% glycerol, and stored at -80°C until use.

### FP antibody binding assessment

To quantify the binding between BG505 or its mutants and FP directed antibodies, the interaction was assessed by BLI using a ForteBio Octet HTX. IgG antibodies were loaded onto Anti-human Fc (AHC) tips at 20µg/ml for 3 minutes, followed by a second baseline phase in PBS for 1 minute. The tips were then dipped into wells with the HIV-1 trimers at 50µg/ml for 5 minutes, followed by a dissociation phase in PBS for 5 minutes. Binding curves were analyzed and background subtracted using the ForteBio Data Analysis 12.0 software, and corrected response values are reported.

### ACS202 Fab and BG505_RnS_7mut_A517E trimer preparation

Antibody ACS202 IgG was purified as described above. Briefly, the codon-optimized variable domains of heavy chain and light chains were cloned into a VRC8400 vector as previously described ^30^, with a HRV3C cleavage site on the heavy chain hinge region. The antibody was expressed by transient transfection in Expi293 cells with plasmids encoding heavy-and light-chain genes. The antibody IgG was purified from Protein A column and then cleaved by HRV3C protease to obtain the antibody Fab, which was purified by passing through a Protein A column and a Superdex 200 column with PBS buffer. The BG505_RnS_7mut_A517E trimer for cryo-EM sample preparation was produced as described above.

### Cryo-EM structural analysis

The ACS202 Fab was mixed with BG505_RnS_7mut_A517E trimer at ∼4.5 molar ratio (Fab to trimer) at a total protein concentration of ∼5 mg/ml. DDM (stock concentration of 1 mM) was added to a final concentration of 0.1 mM. 2.7 μl of the mixture was pipetted to Quantifoil R 2/2 gold grids with vitrobot Mark IV (Thermo Fisher Scientific) using parameters set to 4 °C with 95% humidity and vitrified with no wait time, 2 s blot time, and blot force of -5. Data were acquired with a 300Kv Titan Krios (Thermo Fisher Scientific) equiped with Gatan BioQuantum K3 and operated in the super-resolution mode (0.415 Å/pixel raw unbinned, 0.83 Å/physical pixel) using serialEM ^31^. Data were processed using cryoSPARC 3.3 ^32^ for patch motion correction, patch CTF estimation, particle picking with Blob picker, 2D classifications, *ab initio* 3D reconstructions, homogenous and non-uniform refinements. The final reconstruction map was calculated with non-uniform refinement with C3 symmetry.

Initial rigid body fits of HIV-1 trimer and Fab to the cryo-EM reconstructed maps were performed with Chimera ^33^ using the BG505 Env trimer structure from PDB: 6vi0 ^34^ and the ACS202 Fab structure from PDB: 6nc2 ^27^ as starting models. The mutations in BG505 Env sequence were manually built with Coot ^35^, and the coordinates were refined to fit the electron density map by an iterative process of manual fitting with Coot ^35^ and real space refinement with Phenix ^36^. Molprobity ^37^ was used to evaluate the structural quality. Figures were generated with Chimera and PyMOL ^38^.

### Molecular dynamics simulations

Molecular dynamics (MD) simulations were performed for energetic analysis of antibody-fusion peptide (FP) complexes and ACS202 bound BG505 complexes of mutants. Ten antibody-FP structures (0PVc.01 WT (PDB:6MQC), 0PVc.01 V513E, VRC34(PDB:5I8E), VRC34 G516M, VRC34 G514S, vFP16.02(PDB:6CDO), vFP16.02 G514S, PGT151(PDB:5FUU), PGT151 V513E, PGT151 A512Y) were obtained and mutated by PyMOL ^38^ with the lowest clashing rotamer. Specifically, PGT151-FP complex was truncated from trimer (PDB:5FUU) and constant region of Fab was modelled by VRC34-FP complex structure. Cryo-EM structures of ACS202 in complex with BG505 Env trimer with 517A and mutant 517E were prepared for simulations by modeling missing residues using YASARA (http://www.yasara.org/) and man-5 glycosylation by CHARMM-GUI Glycan Reader and Modeler ^39^. CHARMM-GUI ^40,41^ was used to generate necessary setups for the MD simulations for ten antibody-FP complexes and two BG505_ACS202 mutants. Initial structures were solvated in a 100Å x 100Å x 100Å padded water box for antibody-FP complexes and 197Å x 197Å x 197Å for BG505_ACS202 mutants.

Both systems were neutralized by adding 150 mM of KCl. NAMD2.13 engine ^42^, with CHARMM36 force field ^43,44^ was used to run MD simulations. The water molecules were represented as TIP3P water parameterization ^45^. The electrostatic interactions under periodic boundary condition were computed using particle-mesh Ewald (PME) summation ^46^ with a grid spacing smaller than 1 Å.. The system was performed energetic minimization by 10,000 conjugate gradient steps and equilibrated using a linear temperature gradient. The temperature of the system went up to 310 K in 125,000 steps. The length of hydrogen bonds was constrained with the SHAKE algorithm ^47^ with 2 fs time step. The production step of MD was conducted for 200 ns with NPT (1.01325 bar, 300K) ensemble by Nosé-Hoover Langevin barostat ^48,49^ and Langevin thermostat ^50^ with a damping coefficient 1 ps-1. The final trajectories were used to analyze residue pair energy analysis.

### Residue pair energy calculation

Pairwise residue energy analysis was performed by gRINN, ^51^ for representative frames of each cluster after *Cα* principal component analysis of MD trajectories and clustering on the 2D projections using the meanshift algorithm.

### Statistical analysis

Statistical analysis was performed using custom python scripts and the scipy.stats package. All the code to process data and generate figures will be publicly available.

## Competing interests

The authors declare no competing interests.

## Notes

https://www.ebi.ac.uk/emdb/EMD-41461

### Competing Interest Statement

The authors have declared no competing interest.

