## Supplemental Material for "Vaccine-elicited and naturally elicited antibodies differ in their recognition of the HIV-1 fusion peptide"

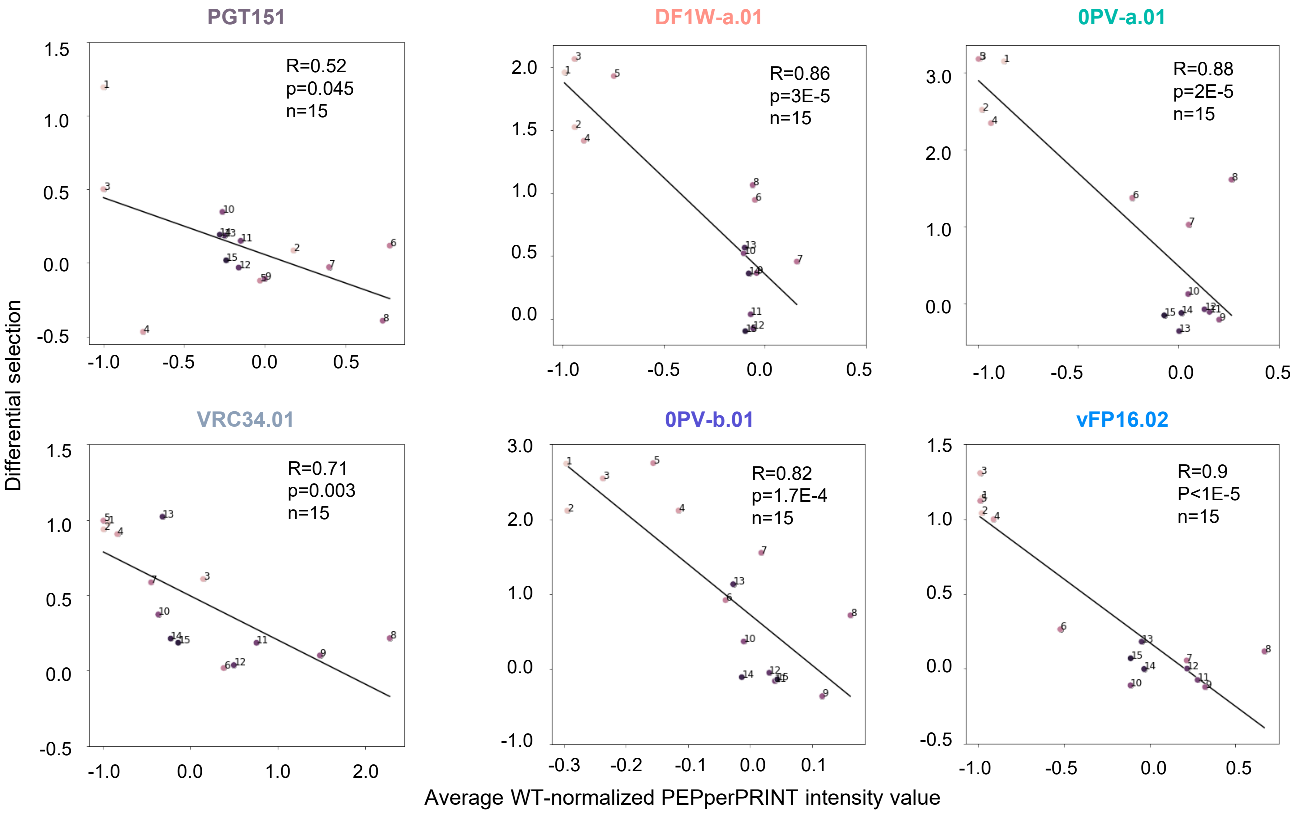


**Figure S1. Peptide-substitution analysis correlates with differential selection measures from deep mutational scanning**

PEPperPRINT WT-normalized intensity values correlate with differential selection values, averaged across site for naturally and vaccine elicited antibodies. Numbers correspond to FP position with 1 corresponding to position 512.

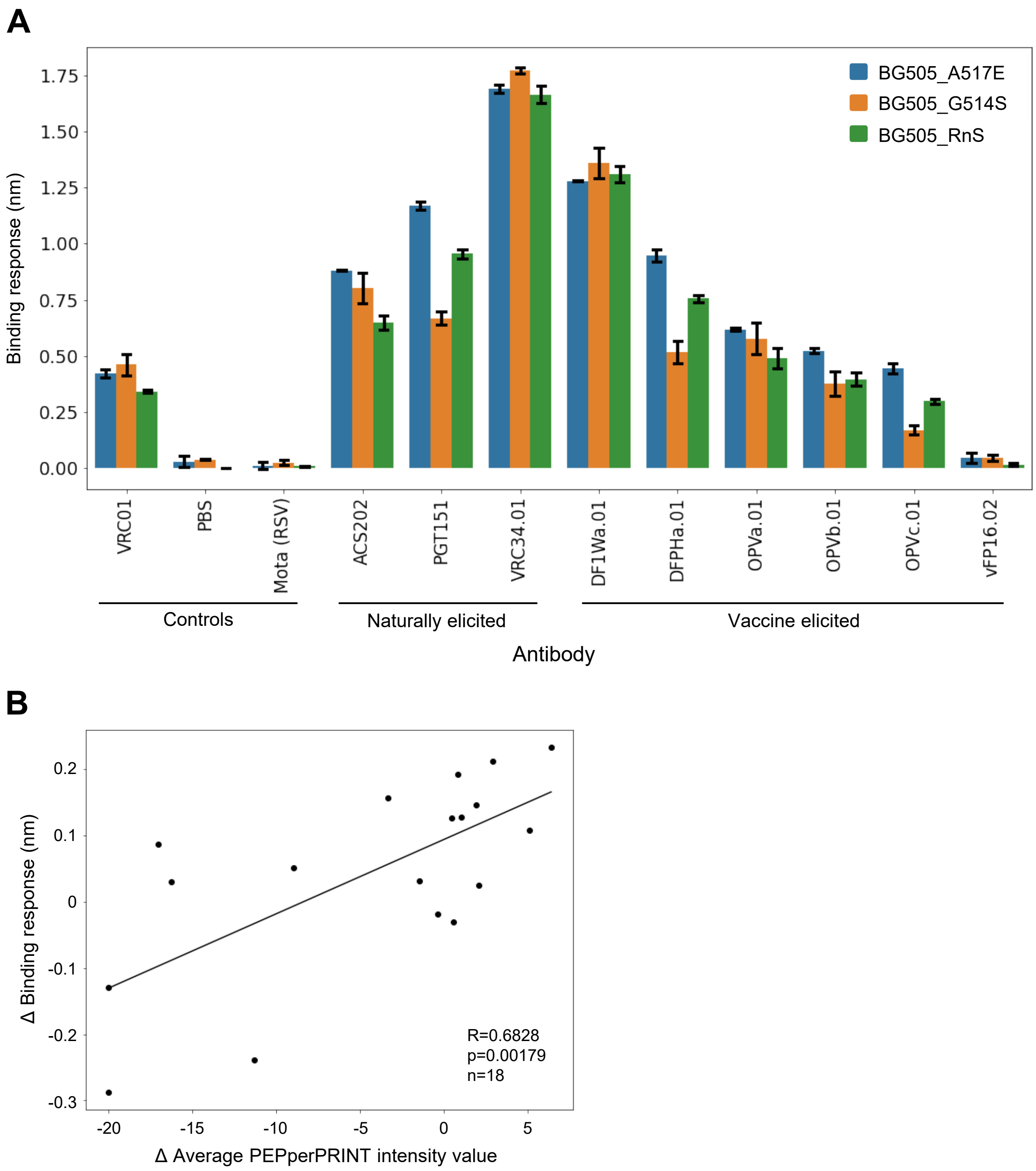


**Figure S2. Octet binding in trimer context correlates with PEPperPRINT binding in peptide context**

(A) Octet binding data for selected antibodies for WT and two mutations (G514S, A517E) with high PEPperPRINT intensity values.

(B) Correlation between octet binding data in trimer context and PEPperPRINT intensity values normalized by mean value per position.

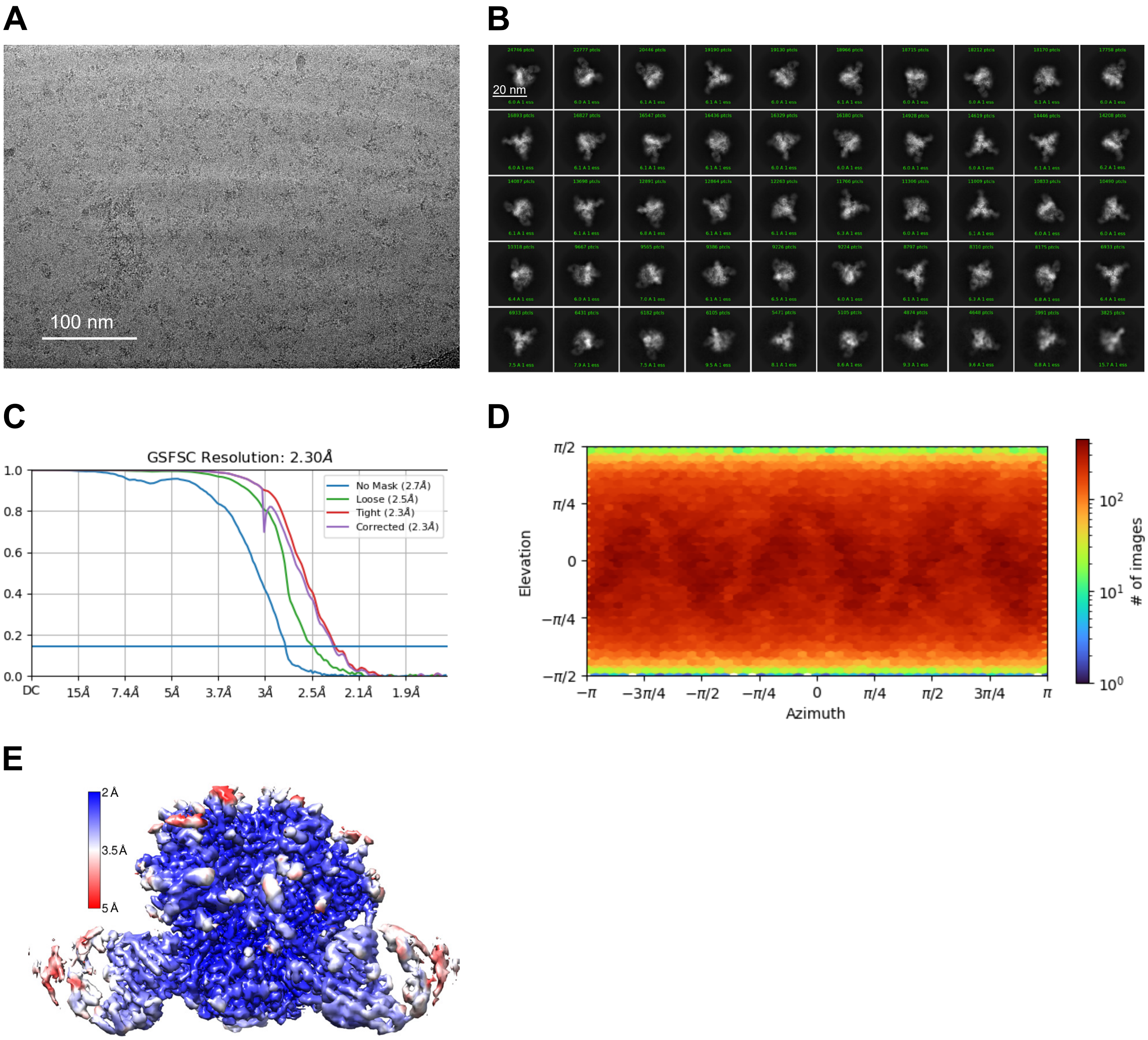


**Figure S3. Cryo-EM validation for ACS202 in complex with BG505_RnS_7mut_A517E, related to Figure 4**

(A) Representative micrograph.

(B) Selected 2D classes.

(C) Fourier shell correlation (FSC) curves showing gold-standard FSC resolution at 2.30 Å.

(D) Orientations of all particles used in the final refinement shown as a heatmap.

(E) 3D reconstruction density map at 2.3 Å from non-uniform refinement with C3 symmetry, with local resolution indicated by color gradient.

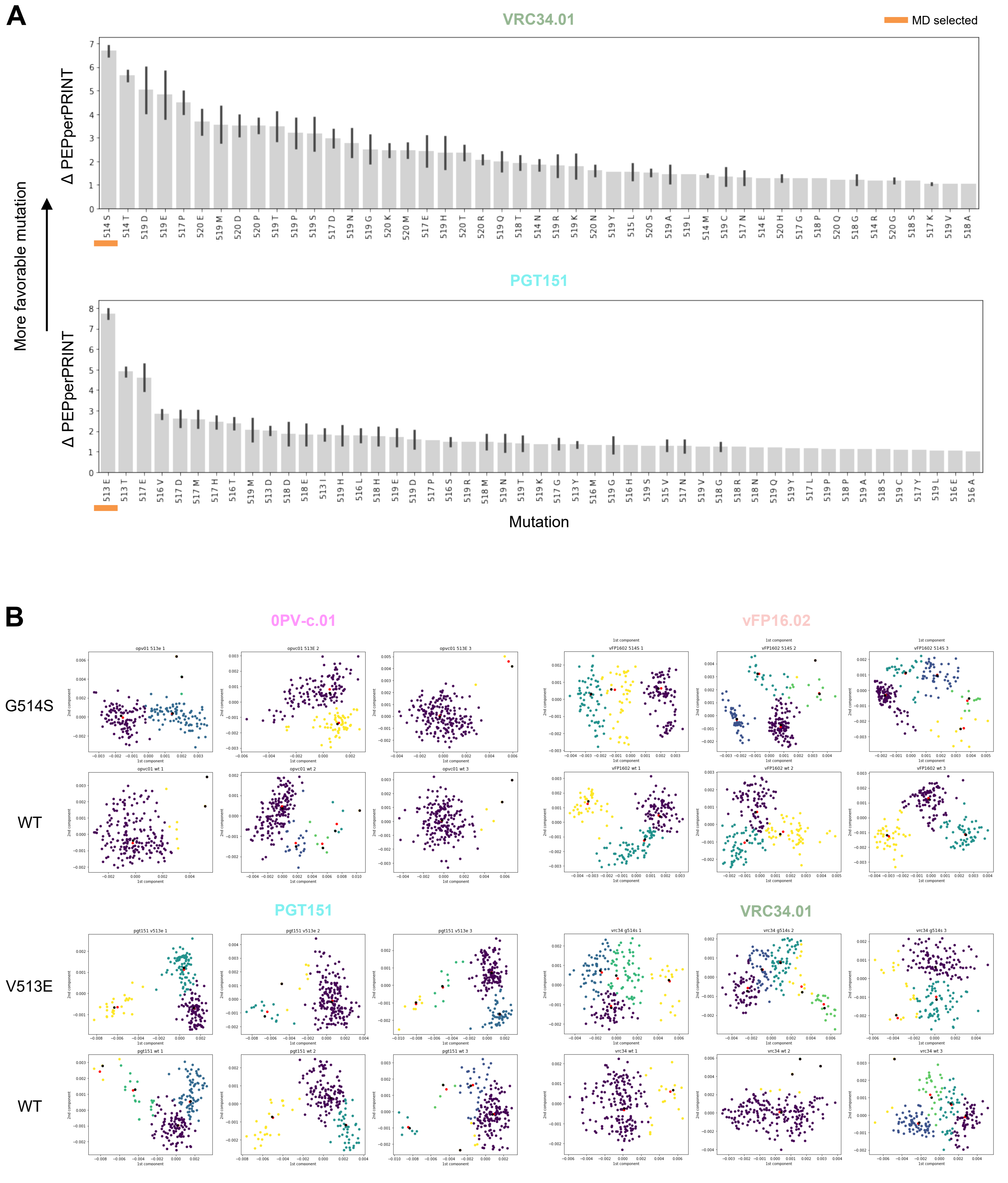


**Figure S4. MD analysis pre-processing, related to figure 5**

(A) Mutant selection for MD simulations by computing the highest ΔPEPperPRINT value between a given naturally elicited antibody value and the mean vaccine elicited antibody value.

(B) PCA analysis of the MD simulations frames from C-α and clustering with the mean shift algorithm. The cluster centroid is in red, and the closest frame to the centroid selected for pairwise analysis in black.

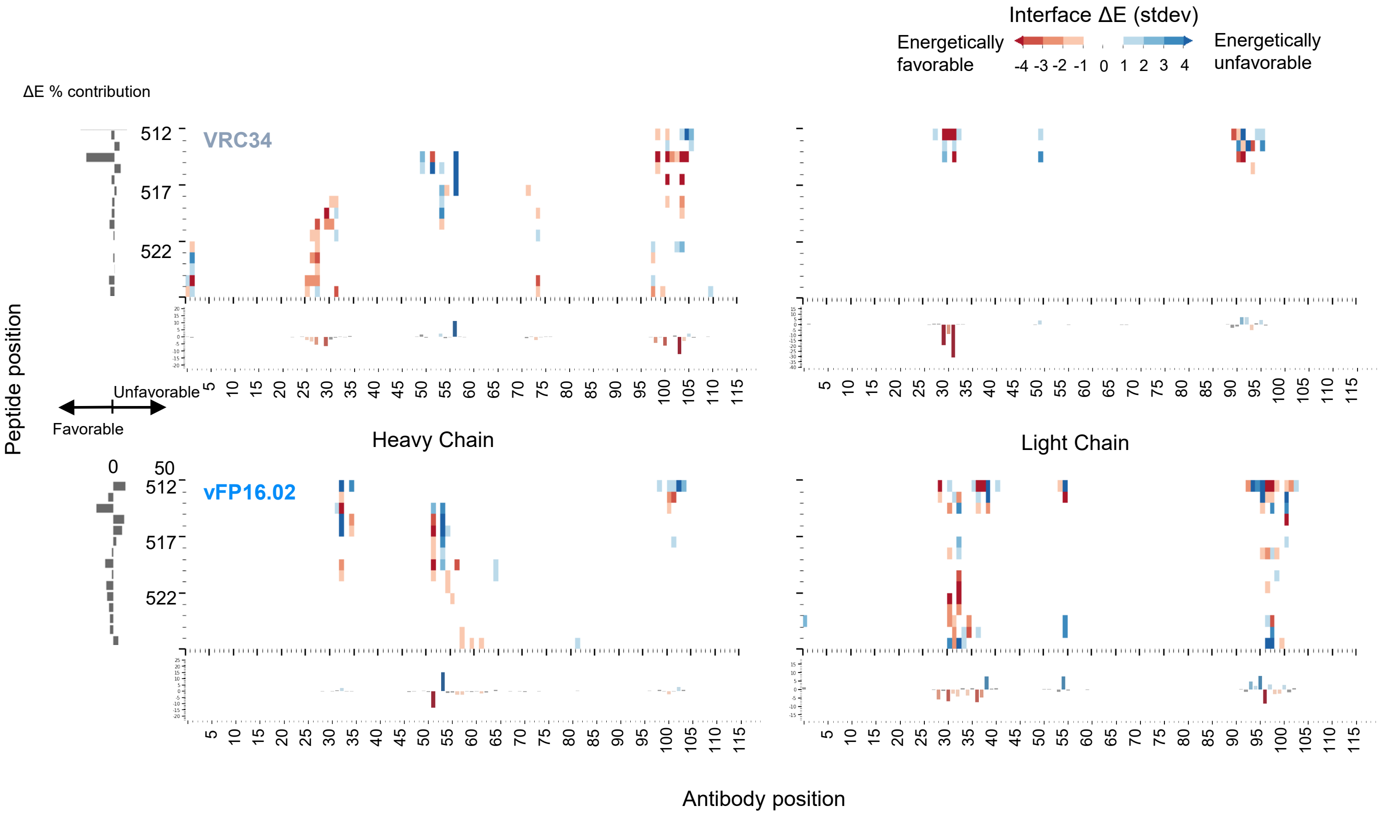


**Figure S5. Atomic level interactions from MD simulation analysis corroborate trends observed by peptide-substitution analysis, related to figure 5**

Pairwise energy analysis reveals specific interactions responsible for the binding differences observed in PEPperPRINT. Numbering of residues is sequential and not in kabat format.

### Table S1. Cryo-EM data and refinement statistics for BG505_RnS_7mut_A517E in complex with ACS202, related to Figure 4

| | |
|---|---|
| **EMDB ID** | EMD-41461 |
| **PDB ID** | 8TOX |
| Data collection | |
| Microscope | FEI Titan Krios |
| Voltage (kV) | 300 |
| Electron dose ($e^-/Å^2$) | 54.40 |
| Detector | Gatan K3 |
| Pixel Size (Å) | 0.415/0.83 |
| Nominal defocus range (μm) | -0.7 to -2.0 |
| Magnification | 105,000 |
| Reconstruction | |
| Software | cryoSPARC v3.3.1 |
| Particles | 635,332 |
| Symmetry | C3 |
| Resolution (Å) ($FSC_{0.143}$) | 2.3 |
| Refinement | |
| Software | Phenix 1.20 |
| Protein residues | 2490 |
| Water | 369 |
| Ligands | BMA: 18; NAG: 114; MAN: 27 |
| CC (box) | 0.82 |
| CC (mask) | 0.85 |
| R.m.s. deviations | |
| Bond lengths (Å) | 0.003 |
| Bond angles (°) | 0.503 |
| Validation | |
| Molprobity score | 1.31 |
| Clash score | 3.52 |
| Rotamer outliers (%) | 0.51 |
| Ramachandran | |
| Favored regions (%) | 97.06 |
| Allowed regions (%) | 2.94 |
| Disallowed regions (%) | 0 |
